# The molecular logic of Nanog-induced self-renewal

**DOI:** 10.1101/374371

**Authors:** Victor Heurtier, Nick Owens, Inma Gonzalez, Florian Mueller, Caroline Proux, Damien Mornico, Philippe Clerc, Agnès Dubois, Pablo Navarro

## Abstract

Transcription factor networks, together with histone modifications and signalling pathways, underlie the establishment and maintenance of gene regulatory architectures associated with the molecular identity of each cell type. However, how master transcription factors individually impact the epigenomic landscape and orchestrate the behaviour of regulatory networks under different environmental constraints is only very partially understood. Here, we show that the transcription factor Nanog deploys multiple distinct mechanisms to enhance embryonic stem cell self-renewal. In the presence of LIF, which fosters self-renewal, Nanog rewires the pluripotency network by promoting chromatin accessibility and binding of other pluripotency factors to thousands of enhancers. In the absence of LIF, Nanog blocks differentiation by sustaining H3K27me3, a repressive histone mark, at developmental regulators. Among those, we show that the repression of Otx2 plays a preponderant role. Our results underscore the versatility of master transcription factors, such as Nanog, to globally influence gene regulation during developmental processes.

## Introduction

Gene regulatory networks driven by master transcription factors (TFs) play pivotal roles over a large spectrum of biological processes, from adaptive cell responses **(Shinozaki et al., 2003)** to cell fate specification during development **(Davidson et al., 2002)**. The key properties of TF networks, shared among cell types, developmental contexts and organisms **(Huang et al., 2005)**, are exemplified by the pluripotency network, which plays a dominant role during early mammalian embryogenesis **(Frum and Ralston, 2015)**. The robustness of this network allows the capture *ex vivo* of the transient biological identity of the pluripotent epiblast through the derivation of self-renewing Embryonic Stem (ES) cells **(Parfitt and Shen, 2014)**, which have enabled identification of key TFs (e.g. Oct4, Sox2, Nanog and Esrrb). The study of processes driving the balance between ES cell self-renewal and differentiation has provided us with a canonical picture of how TF networks operate, establishing self-sustaining regulatory loops and acting together through multiple promoters and enhancers **(Boyer et al., 2005; Loh et al., 2006; Chen et al., 2008; Kim et al., 2008)**. For instance, Oct4, without which pluripotent cells cannot be maintained **(Niwa et al., 2000)**, acts with the TF Sox2 to recognise and bind chimeric motifs **(Chew et al., 2005)** found at a large number of regulatory elements driving ES cell-specific transcription. Oct4 and Sox2 tend also to bind with other TFs, including Nanog and Esrrb, at multiple enhancers across the genome, to combinatorially coregulate a large number of targets. This simultaneous and concerted action over hundreds of common targets ensures extensive redundancy, and, therefore, robust genome-wide responses. How these TFs synergise at or compete for common regulatory elements, and how by these means they individually contribute to the network’s activity, is however not well understood. Moreover, several TFs of the pluripotency network are directly connected to cell signalling, enabling ES cells to establish appropriate responses that are instructed extrinsically. A prominent example is provided by the LIF cytokine, which promotes self-renewal by activating several pluripotency TFs such as Esrrb **(Niwa et al., 2009; Huang et al., 2018)**. Hence, a key function of the pluripotency network is to integrate signalling cues to appropriately respond to changes in the environment, conferring the responsiveness of ES cells and their capacity to readily differentiate. In this regard, it is noteworthy that *Nanog* was first identified as a factor capable of bypassing the requirements for LIF: in the presence of ectopic Nanog expression, ES cell self-renewal is strongly enhanced and completely independent of LIF **(Chambers et al., 2003)**. In the current model, Nanog achieves LIF-independent self-renewal by activating LIF-responsive genes, in particular *Esrrb*. Hence, the Nanog-Esrrb axis and its intersection with LIF signalling represents a major mechanism by which intrinsic and extrinsic cues fine-tune self-renewal and avoid differentiation **(Festuccia et al., 2012)**. Yet, the precise mechanisms by which Nanog, and more generally the pluripotency network, controls differentiation genes are not fully understood. It is known, however, that differentiation genes adopt a particular chromatin state known as “bivalent” **(Azuara et al., 2006; Bernstein et al., 2006)**: while their promoters are enriched for H3K4me3, a mark of gene activity, they are simultaneously embedded within larger domains of H3K27me3, a repressive mark. During differentiation, this state is resolved in either H3K27 or K4me3 in a lineage-specific manner **(Mikkelsen et al., 2007)**. In agreement, Polycomb Group proteins triggering H3K27me3 ensure appropriate cell fate changes **(Boyer et al., 2006; Pasini et al., 2007; Leeb et al., 2010)**. This underscores the importance of H3K27me3 as cells dismantle the pluripotency network, inhibit self-renewal and exit from pluripotency. Whether bivalent chromatin marks are governed by pluripotency TFs remains to be thoroughly addressed.

In this study, we explore the function of Nanog in mouse ES cells using inducible approaches of gain-and loss-of-function. We show that Nanog drives the recruitment of Oct4, Sox2 and Esrrb at thousands of regulatory regions, from where it mainly activates transcription. At these sites, Nanog also recruits Brg1 and promotes chromatin accessibility. On the contrary, to repress transcription Nanog does not recruit these TFs; rather, it frequently inhibits Oct4 or Sox2 binding. Nanog also binds at other enhancers where it acts redundantly with other TFs. However, in the absence of LIF the action of Nanog over these regulatory elements becomes dominant, particularly to promote transcription. This results in Nanog having an expanded action in the absence of LIF. Yet, its expanded repressive activity is not associated with ES cell enhancers. Rather, Nanog is required to maintain H3K27me3 at differentiation-associated genes. This is the case of the TF Otx2, whose downregulation by Nanog leads to LIF-independent self-renewal even when Esrrb is not expressed. Hence, Nanog deploys distinct molecular means to promote self-renewal and counteract differentiation: when the network is fully operative (in the presence of LIF), Nanog rewires its activity; when it is partially dismantled (in the absence of LIF), Nanog represses differentiation genes via H3K27me3. Overall, we reveal different modes and the varied logic employed by Nanog to orchestrate the three main features associated with self-renewal: the inter-dependencies between pluripotency TFs, LIF signalling, and bivalent chromatin domains.

## Results

### Inducible CRISPR-ON ES cells efficiently activate Nanog transcription

The SunTag system was developed as a versatile tool to either visualise specific molecules in live cells or to perform epigenome editing of endogenous loci when coupled to an enzymatically inert dCas9 (**Tanenbaum et al., 2014**). It involves the expression of diffusible antibodies (scFv) that interact with high affinity with 10 copies of the GCN4 epitope linked to an enzymatically inert Cas9 (dCas9). These scFv antibodies are fused to GFP and the potent activator VP64, such that upon expression of a gRNA targeting a given genomic region, several VP64 molecules are brought about with high efficiency and specificity. To provide increased flexibility to the system, and facilitate the generation of cell lines carrying an inducible CRISPR-ON system, we engineered a single vector expressing the two SunTag moieties under the control of a Tetracycline Responsive Element. Moreover, dCas9 is linked to BFP and HpH through P2A and IRES sequences, respectively (Fig. S1A). Hence, upon induction of the system with Doxycycline (Dox), the cells are expected to become green, blue, and Hygromycin-resistant, providing a high tractability. This vector was introduced in ES cells together with the rtTA activator: two clones (C1 and C2) showing a high percentage of green/blue cells upon Dox treatment and a strong induction of dCas9 and VP64 (Fig. S1B, C), were selected. They both self-renew normally and differentiate in the absence of LIF; their karyotypes are also normal (Fig. S1D).

Next, we introduced to C1 and C2 a vector expressing a gRNA targeting the minimal Nanog promoter and validated binding of dCas9/VP64 with good specificity and inducibility (Fig. S1E). This was accompanied by increased histone H3 acetylation around the promoter (Fig. 1A, B), as expected given the ability of VP64 to recruit histone acetyl-transferases (**Utley et al., 1998**), in the context of presumably unaltered nucleosomal organisation as evaluated by total H3 analysis (Fig. 1B). We also assessed *Nanog* expression over the course of 6 days of induction, and showed that both Nanog pre-and mRNA were induced from day 1 onwards (Fig. 1C), leading to an increase of Nanog protein levels (Fig. S1C). We also found that the increase of *Nanog* expression was due both to stronger and more frequent transcriptional bursts (Fig. 1D and Fig. S1F, G). Finally, we analysed the effects of Dox administration at the proximal −5kb enhancer of *Nanog*: upon induction, we found both sense and anti-sense enhancer transcription to be increased (Fig. 1E). Whether this is due to the proximity of these two regulatory elements or to a functional influence of the promoter on the enhancer, remains to be determined. In conclusion, we have generated Dox-inducible SunTag ES cells to activate endogenous promoters and dissect the subsequent consequences.

**Fig. 1.**
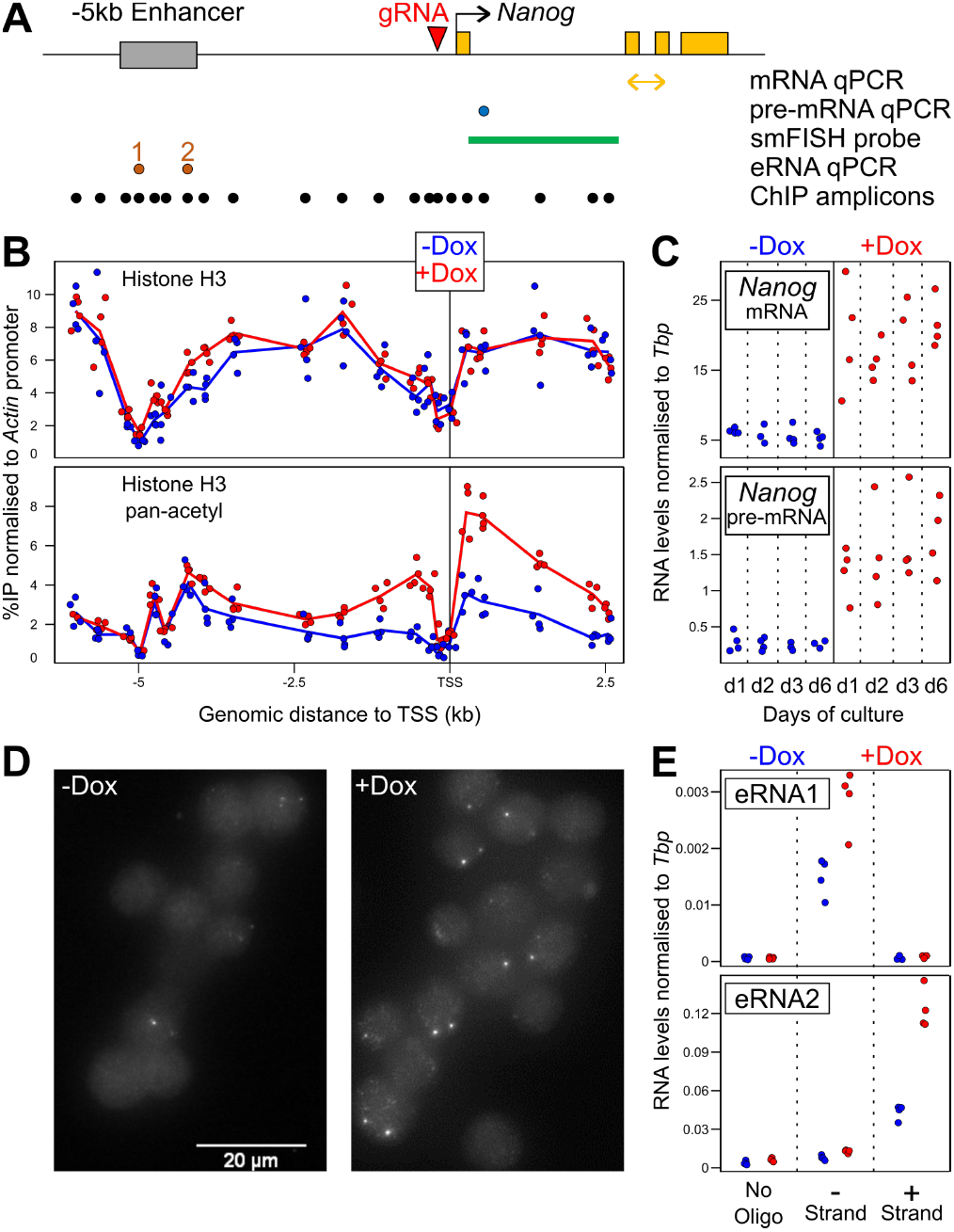
CRISPR-ON ES cells for Dox-inducible activation of endogenous *Nanog*. **(A)** Schematic representation of the *Nanog* locus (black arrow: promoter; yellow boxes: exons; grey box: *Nanog* enhancer; red arrowhead: gRNA). Below, the position of the amplicons and probe used for the assays indicated on the right is shown. **(B)** ChIP across the *Nanog* locus monitoring total histone H3 (top) and pan-acetyl H3 (bottom) in the absence (blue) and after 72h of Dox treatment (red). Each dot represents normalised %IP measured in individual replicates and lines the averages. **(C)** Normalised levels of Nanog mRNA (top) and pre-mRNA (bottom) after the indicated number of days in the absence (blue) and the presence (red) of Dox. Each dot represents measurements in individual replicates as in (B). **(D)** Representative smFISH image using intronic *Nanog* probes before and after 72h of Dox induction. **(E)** Normalised levels of eRNA production from the −5kb enhancer presented as in (C). In all panels, n=4; 2 with each independent SunTag clone.

### Definition of Nanog responsive genes

Upon Dox induction of *Nanog* in our SunTag cells we observed a 2-fold increase of Nanog binding to a panel of regulatory elements displaying a wide range of enrichment levels (Fig. 2A). This suggests that Dox induction may lead to functional consequences. However, the two main targets of Nanog that have been previously identified, *Esrrb* and *Klf4* (**Festuccia et al., 2012**), did not show any variation in expression levels over the course of 6 days of endogenous *Nanog* induction (Fig. 2B). Prompted by this unexpected observation, we performed RNA-seq to comprehensively study the global response to Dox treatment. We found a small number (163) of transcripts that were either up-or down-regulated (Fig. 2C top); neither *Klf4* nor *Esrrb* were among the induced genes (Table S1 and Fig. S2A, B). Nevertheless, the vast majority of genes that have been previously identified as responding to Nanog levels (**Nishiyama et al., 2009; Festuccia et al., 2012**), do exhibit the appropriate expression changes in our SunTag cells (Fig. S2C). To further validate our list of Nanog-responsive genes, we performed a complementary analysis using previously established *Nanog*-null cells (44iN) expressing a Dox-inducible *Nanog* transgene (**Festuccia et al., 2012**). The cells were grown in the continuous presence of Dox, which was then removed for 24 hours leading to a nearly complete loss of *Nanog* expression (Fig. 2C bottom). The number of responsive genes (141) observed with this strategy was also small (Fig. 2C and Table S1); they intersected with excellent statistical significance with the genes identified in the Sun-Tag cells (p<1e-53). Moreover, we found the expression of genes significantly regulated in only one system, to nevertheless display highly coherent expression changes in the other system (Fig. 2D). Hence, to improve statistical power and expand on Nanog targets we combined the SunTag and 44iN datasets to test for those genes with coherent Nanog response across both systems (Fig. 2E and Fig. S2D). Combining with those genes already identified, this resulted in 457 genes (Fig. 2E and Fig. S2D), which generally display extremes of expression differences between Dox-treated SunTag (high *Nanog*) and untreated 44iN cells (low/absent *Nanog*); they globally behave in a concordant way when long-term *Nanog*-null cells are compared to wild-type cells (Fig. S2D), or when their expression is analysed in published datasets (Fig. S2E). Genes activated by Nanog are enriched in regulators of stem cell maintenance (FDR<2.99e-16), while repressed genes are enriched in differentiation processes such as nervous system development (FDR<6.89e-10). Hence, we have defined the compendium of Nanog-responsive genes in undifferentiated ES cells with unprecedented completion. Strikingly, neither *Klf4* nor *Esrrb* belong to our list of Nanog-responsive genes.

**Fig. 2.**
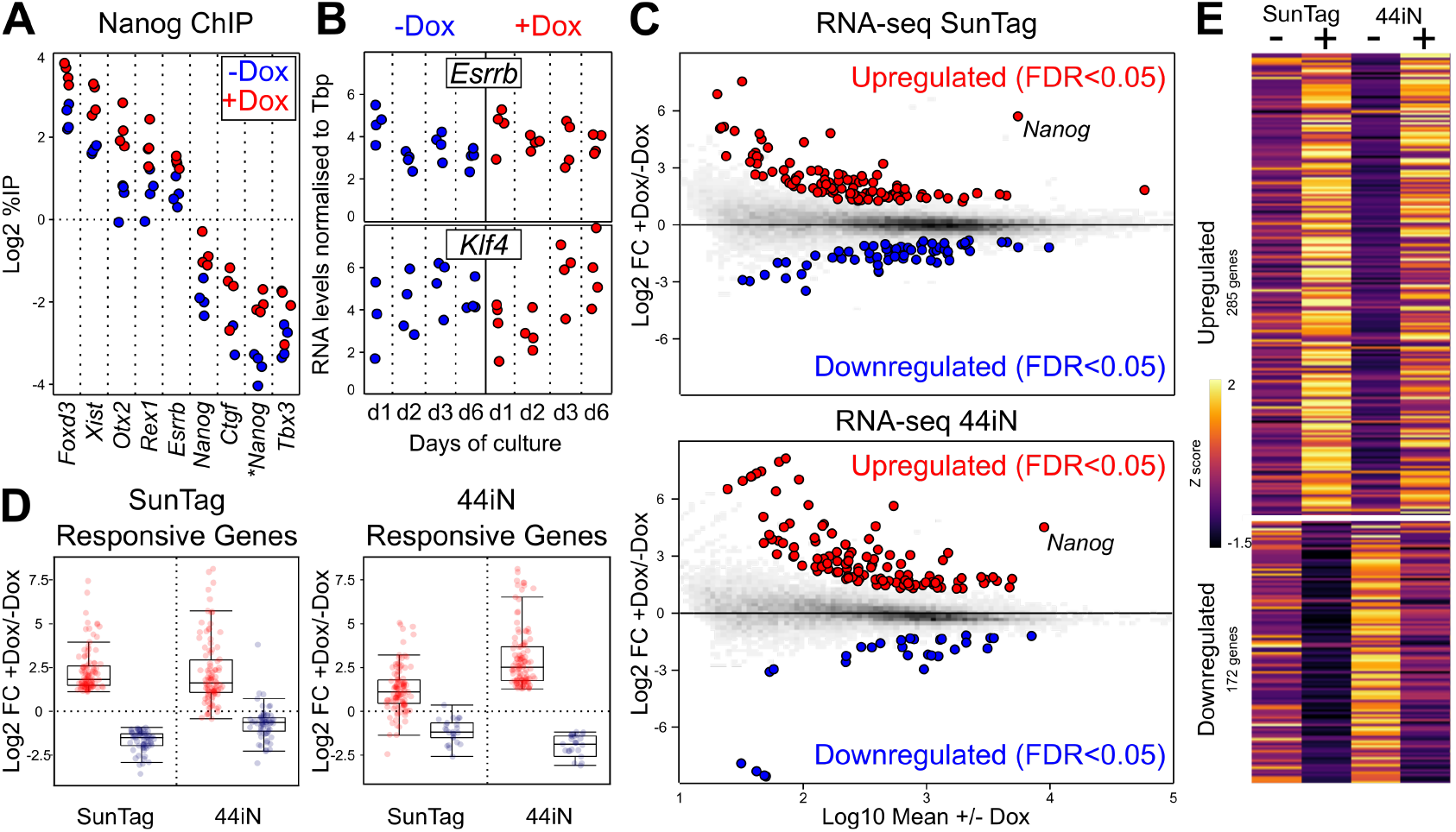
Identification of Nanog-responsive genes. **(A)** ChIP analysis of Nanog binding across a set of targets, as indicated on the X-axis. Each dot represents measurements in individual replicates (n=4; 2 for each independent SunTag clone) **(B)** Normalised levels of *Esrrb* (top) and *Klf4* (bottom) mRNA after the indicated number of days in the absence/presence of Dox. Each dot represents measurements in individual replicates as in (A). **(C)** MA Plots displaying log2 fold changes as indicated on the Y-axis as a function of average expression. RNA-seq was performed in both SunTag (72h Dox induction) and 44iN cells (24h Dox withdrawal). Red and blue represent differentially expressed genes (FDR<0.05). **(D)** Boxplot of the log2 fold change for the genes identified in (C) as upregulated (red) or downregulated (blue) by Nanog, in either SunTag (left) or 44iN cells (right), measured in both inducible systems as indicated on the X-axis. **(E)** Heat map representing gene expression z scores of all transcripts identified by combining SunTag (FDR < 0.05), 44iN (FDR <0.05) and SunTag/44iN likelihood ratio test (FDR < 0.05) datasets.

### Nanog rewires the pluripotency network to control its targets

Having established a list of Nanog-responsive genes, we aimed at exploring the mechanisms by which Nanog influences their expression, focusing on a potential role of Nanog in modulating binding of other regulators such as pluripotency TFs (Oct4, Sox2 and Esrrb) and the chromatin remodeller Brg1 that is functionally associated with self-renewal (**Ho et al., 2009; Kidder et al., 2009**). To do this, we first established a list of 27,782 regulatory elements bound by Nanog using six datasets derived from four independent published studies (Table S2) and used 44iN cells to address how Nanog impacts TF binding and chromatin accessibility at these sites. We noticed that at some Nanog binding regions, Esrrb, Oct4, Sox2, and Brg1, display a strong reduction of binding and decreased chromatin accessibility, after 24 hours of Dox withdrawal (Fig. 3A). This observation can be generalised to a large proportion of regions and is particularly prominent in the case of Esrrb (Fig. 3B). We then divided Nanog binding regions in two major groups Fig. 3B and Fig. S3, based on the presence of other TFs (regions of co-binding) or not (Nanog-solo regions). In Nanog-solo regions, which display lower levels of Nanog binding, the chromatin is less accessible irrespective of Nanog (Fig. 3B and Fig. S3), indicating that additional factors may be recruited. Strikingly, when we computed the number of Nanog-responsive genes as a function of the distance to Nanog binding regions, we observed that both activated and downregulated genes are particularly enriched in the vicinity of co-binding regions and not of Nanog-solos (Fig. 3C). Moreover, while activated genes tend to be located distally (within 10 to 100kb), downregulated genes also show a significant enrichment over closer distances (<10kb). To further explore the relationships between Nanog and other TFs, we used k-means clustering to identify 8 subgroups of Nanog binding sites (Fig. 3D and Fig. S3). In the first 4 clusters, the depletion of Nanog leads to an acute loss of TF binding; collectively these regions are strongly associated with the activation of Nanog targets (Fig. 3C). In contrast, clusters 5-8 are significantly associated with genes repressed by Nanog and the effects of its depletion are more nuanced (Fig. 3C, D). More specifically, clusters 1 to 3 display a nearly total loss of TF binding in the absence of Nanog, along with a marked decrease in chromatin accessibility (Fig. 3D and Fig. S3). These 3 clusters, in particular clusters 1 and 2, are associated with genes activated by Nanog (Fig. 3C). At cluster 4, however, chromatin accessibility shows minimal variations and, while very strong Oct4 binding is nearly completely lost upon Nanog depletion, Sox2 is not affected. Since Brg1 is particularly low across this cluster, Sox2 may recruit other chromatin remodellers to render the chromatin accessible at these regions, in a Nanog-independent manner. Accordingly, the correlation with Nanog-responsive genes of cluster 4 is weaker (Fig. 3C). Overall, at more than 6000 regions (clusters 1 to 3), Nanog plays a chief role in establishing functional and accessible regulatory regions capable of recruiting different combinations of TFs to activate its targets. Con-versely, at clusters 5 to 8, the effects of the loss of Nanog are rather small both at the level of TF binding and of chromatin accessibility (Fig. 3D and Fig. S3). This suggests that Nanog-mediated repression uses radically different mechanisms, which are not based on the increased recruitment of Esrrb, Oct4 and Sox2. Rather, clusters 7 and 8 display increased Oct4 and Sox2 binding in the absence of Nanog, respectively, suggesting that Nanog downregulates the genes functionally linked to these two clusters by blocking Oct4 or Sox2 recruitment (Fig. 3D and Fig. S3). At other enhancers associated with genes repressed by Nanog, showing no alteration of Oct4 and Sox2 occupancy (clusters 5 and 6), Nanog may block the otherwise activatory function of other TFs. In conclusion, Nanog wires the pluripotency network by fostering TF recruitment and chromatin accessibility at distal regulatory elements to act as an activator, and uses different mechanisms, including the impairment of Oct4/Sox2 recruitment, both at promoter-proximal and distal regulatory elements of the genes it represses.

**Fig. 3.**
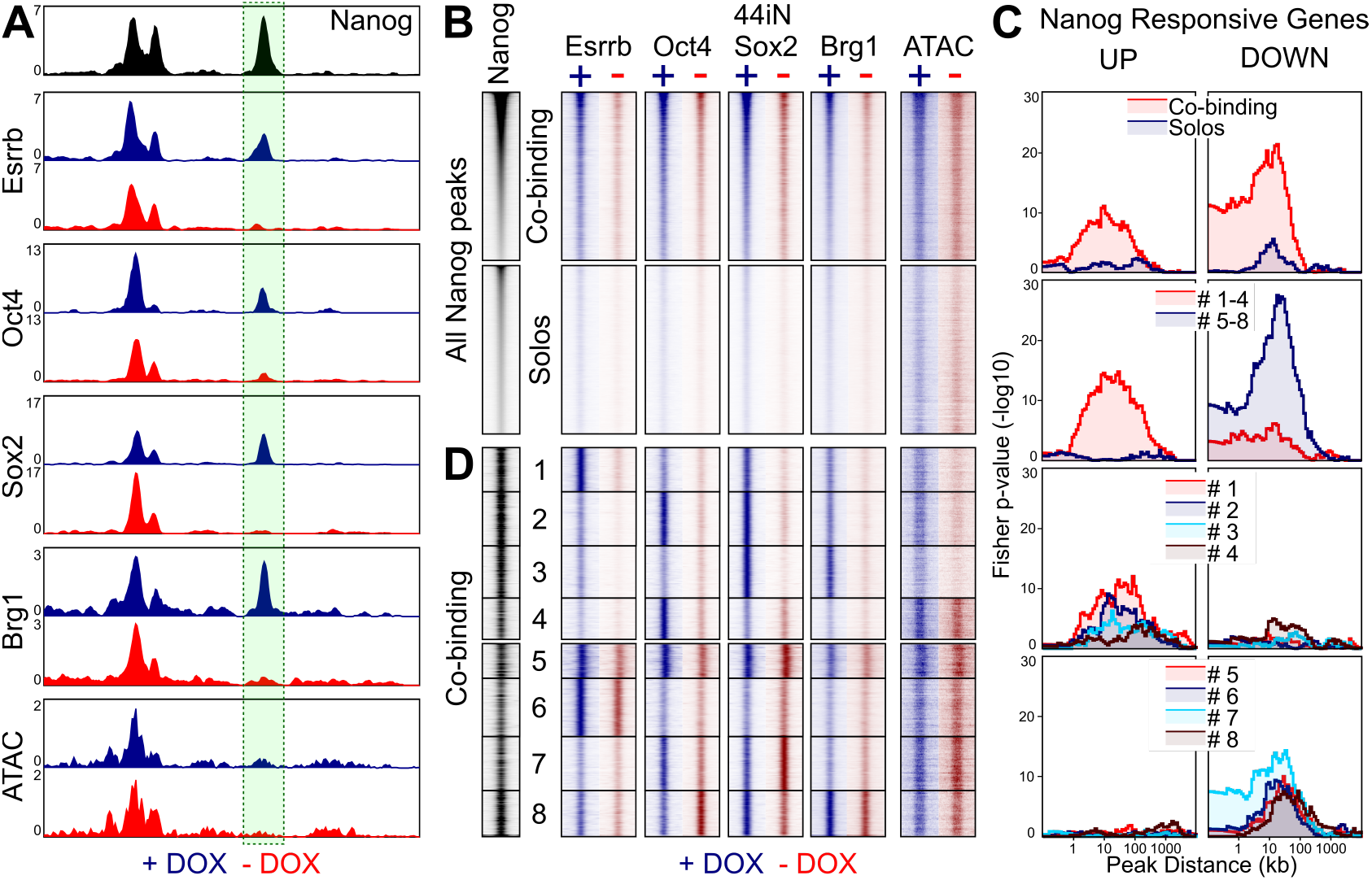
Nanog rewires the pluripotency network. **(A)** Representative enrichment profiles (reads per million) of the indicated TFs (ChIP-seq) and chromatin accessibility (ATAC-seq) across an 8kb-long genomic region (mm9: chr10:33,952,000-33,960,000). In black, average signal from 6 publicly available Nanog datasets. In blue and red, average signal of 44iN cells cultured in the presence or absence of Dox, respectively. The peak highlighted by a green box shows reduced signal in the absence of Dox; note an upstream peak displays increased Sox2 binding. **(B)** Heat map of average enrichment levels in each indicated condition (+/-Dox) across 0.5kb centred on the Nanog peak summit at all Nanog sites (27,782), ranked from high to low Nanog. The regions were split in two groups depending on the presence (co-binding) or absence (Solos) of other TFs. **(C)** Plots displaying the −log10 Fisher p-value (Y-axis) of the enrichment of genes up-(left) or downregulated (right), at a given distance (X-axis) from specific groups of Nanog binding sites identified in (B, D), as indicated in the colour-coded insets. **(D)** Heat map corresponding to 8 clusters identified on the basis of TF co-binding and the effects of Dox withdrawal, presented as in (B) without ranking for Nanog binding levels.

### Nanog confers LIF-independent self-renewal in the absence of *Esrrb* induction

The strong influence of Nanog on the efficiency of self-renewal has been proposed to be largely mediated by Esrrb (Festuccia et al., 2012). Therefore, we were not expecting our SunTag cells endogenously activating *Nanog* to exhibit increased self-renewal capacity, given that *Esrrb* and other genes involved in self-renewal, such as *Klf4* (Niwa et al., 2009), are not strongly induced (Fig. 2B). To test this, we initially plated our SunTag lines at clonal density together with the parental controls lacking the *Nanog* gRNA, and cultured them in the presence or absence of Dox for 6 days (Fig. 4A). In the presence of LIF, we could not observe any major change in the efficiency of self-renewal. In contrast, in the absence of LIF, when virtually all the colonies display complete or partial signs of differentiation in all controls, cells with enhanced endogenous *Nanog* expression generated a substantial proportion of undifferentiated colonies (Fig. 4A, B). To further validate that these cells are bona-fide ES cells, we harvested them at the end of the clonal assay and performed two complementary assays. First, we re-plated them in 2i medium lacking serum (**Ying et al., 2008**), where only truly undifferentiated cells proliferate: both clones gave rise to typical spherical and undifferentiated colonies (Fig. 4C). Second, we re-plated them at clonal density in the absence of LIF and the presence/absence of Dox: only in the presence of Dox did we recover undifferentiated colonies; in the absence, all the cells differentiated (see below). This demonstrates that the exposure to Dox does not alter the differentiation capacity of our cell lines upon its withdrawal. We conclude, therefore, that Dox-induction of *Nanog* confers to our SunTag lines the ability to self-renew in the absence of LIF, a definitive proof of the efficiency of our CRISPR-ON strategy to study Nanog function. Strikingly, LIF-independent self-renewal was attained in the absence of any apparent induction of Klf4 and Esrrb mRNAs (Fig. S4A) or proteins (Fig. 4D). Therefore, to explore both the magnitude of the differentiation blockade at the molecular level, and to identify potential Klf4/Esrrb-independent mechanisms underlying Nanog-mediated self-renewal, we performed transcriptomic analyses. In control cells that were not stimulated by Dox, a large number of genes responded to LIF withdrawal (>5000) and exhibited important quantitative differences (Fig. S4B, C, D). In the presence of Dox, the magnitude of the expression changes of these LIF-responsive genes was globally diminished (Fig. S4B), even though the vast majority of pluripotency genes remained strongly downregulated (Fig. S4C, D). In fact, not all genes that respond to LIF withdrawal were rescued by *Nanog* induction to the same extent, with only around 20% being efficiently rescued (Fig. 4E, Fig. S4E and Table S1). This argues against the idea that the presence of substantial numbers of undifferentiated cells may explain all the expression changes measured upon Dox induction in the absence of LIF.

**Fig. 4.**
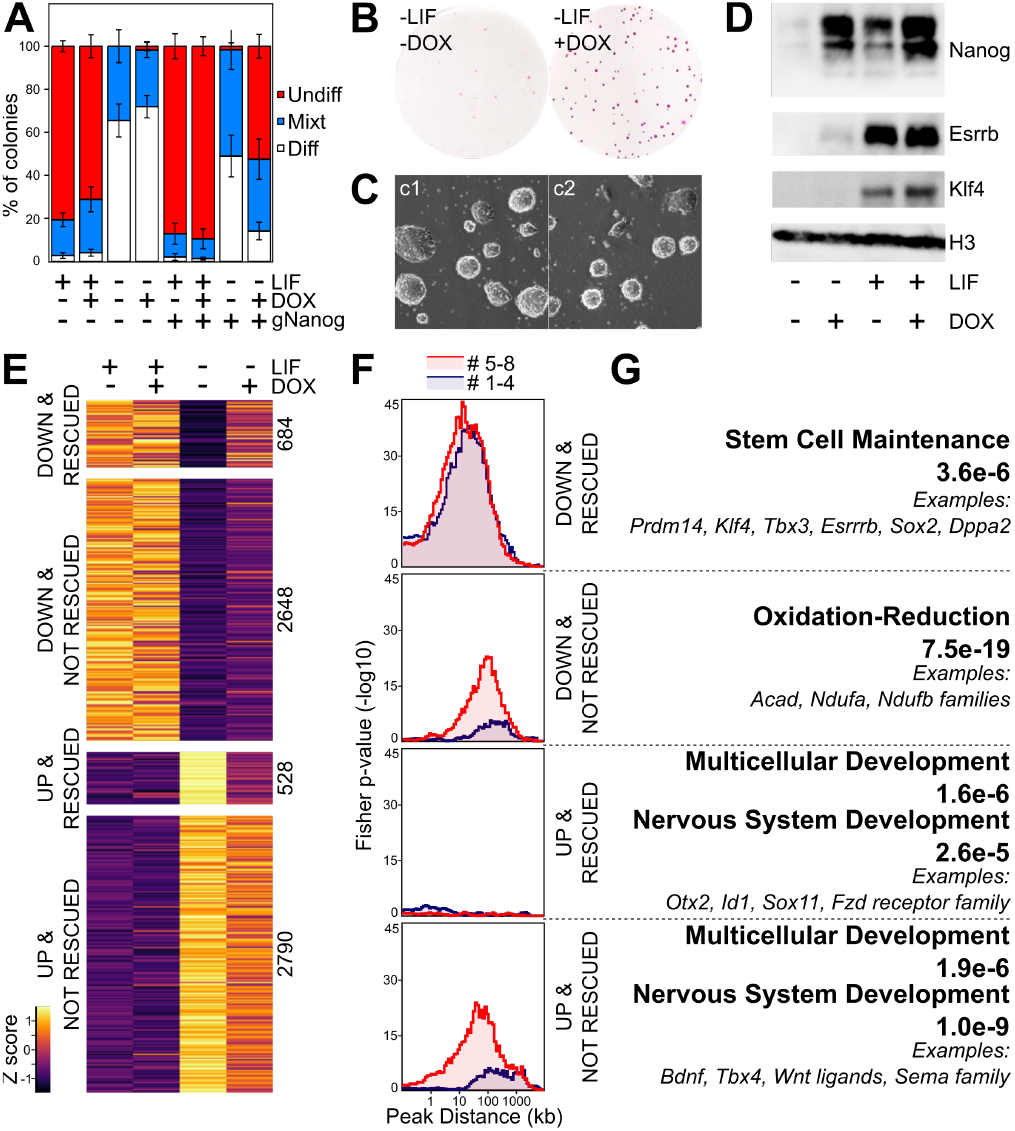
Endogenous induction of *Nanog* mediates LIF-independent self-renewal by non-canonical mechanisms. **(A)** Histogram representing the percentage of undifferentiated (red), mixt (blue) and differentiated (white) colonies (Y-axis) counted after 6 days of clonal growth in the indicated conditions (X-axis). Error bars represent std. dev. (n=4). **(B)** Representative image of Alkaline Phosphatase stained colonies after 6 days of clonal growth in the presence/absence of Dox. **(C)** Representative image of the two SunTag clones after culturing them in 2i for 3 days following a 6-day clonal assay in the absence of LIF and the presence of Dox. **(D)** Representative Western-Blot of Nanog, Esrrb and Klf4 after 3 days of culture in the indicated conditions. **(E)** Heat map representing gene expression z scores across 4 groups of transcripts, as indicated in the left. UP/DOWN refers to their expression changes after 3 days of LIF deprivation (FDR<0.05); rescued versus not rescued indicates whether Nanog significantly alleviates their misregulation (FDR<0.05). **(F)** Cumulative plots displaying the −log10 Fisher p-value (Y-axis) of the enrichment of genes belonging to the groups identified in (E) at a given distance (X-axis) from Nanog binding clusters 1 to 4 (red) and 5 to 8 (blue), as established in Fig. 3. **(G)** Representative Gene Ontology terms enriched in each group of genes. The FDR is indicated together with selected examples.

### Increased regulatory potential of Nanog in the absence of LIF, due to a loss of redundancy

In the absence of LIF, the effects of endogenous Nanog induction are largely maximised: if in the presence of LIF we identified 285 up and 172 downregulated genes, in its absence these numbers raised to 856 and 589, respectively (Table S1). It appears, therefore, that LIF signalling attenuates the relative impact of Nanog on the ES cell transcriptome. To explore this further, we established the associations between the clusters of Nanog binding regions we have identified (Fig. 3), with four groups of genes: genes down-or upregulated upon LIF withdrawal and, among these two categories, those that Nanog can or cannot partially rescue by activating or repressing them, respectively (Fig. 4E). We observe that only one group, constituted of genes repressed by Nanog in the absence of LIF, is not enriched in any Nanog binding region that we have studied (Fig. 4F and Fig. S4F). In contrast, the group of genes downregulated upon LIF withdrawal, and rescued by Nanog, is similarly enriched for Nanog binding regions where Nanog leads the recruitment of other TFs (clusters 1 to 4) than for those where it does not (clusters 5 to 8; Fig. 4F and Fig. S4F). The activatory potential of Nanog through clusters 1 to 4 was already established from the previous analysis. However, the vast majority of genes upregulated by Nanog in the absence of LIF were not activated in LIF containing medium (>700 genes, Table S1). This suggests that in the presence of LIF, these genes are redundantly controlled by other LIF-dependent TFs, either through regions belonging to clusters 5 to 8 or through other regulatory elements where Nanog does not bind. Since clusters 5 to 8 were previously associated with genes repressed by Nanog in the presence of LIF, their enrichment in the vicinity of genes activated by Nanog in the absence of LIF implies that they are constituted by at least two functional categories: enhancers that are blocked by Nanog in the presence of LIF, leading to the downregulation of Nanog targets, and enhancers where Nanog also acts as an activator but redundantly to other LIF-dependent factors, most likely Esrrb. In agreement, this group is strongly enriched in genes of the pluripotency network (Fig. 4G), which are known to be controlled by several pluripotency TFs and from multiple distinct enhancers. The level of upregulation of these Nanog-activated genes is, however, relatively minor and the rescue of pluripotent TFs, including *Esrrb* and *Klf4*, marginal (Fig. 4D and Fig. S4A, C). Finally, LIF-responsive genes that are not rescued by Nanog are associated with clusters 5 to 8 (Fig. 4F). This indicates that a large number of regulatory regions of the pluripotency network for which Nanog has a modest functional impact are present within these clusters. These regions activate or repress genes prior to differentiation in a Nanog-independent manner; upon LIF-withdrawal, their activity is likely invalidated with the ensuing consequences on gene expression even when *Nanog* is induced. These Nanog-independent, LIF-responsive genes, are closely associated with cluster 6 (Fig. S4F), which is dominated by Esrrb (Fig. 3D and Fig. S3), a prominent LIF target. This is particularly true for genes downregulated upon LIF withdrawal; satisfactorily, given the known role of Esrrb as a general regulator of metabolism and energy production (**Sone et al., 2017**), these genes are enriched for related terms such as oxidation-reduction (Fig. 4G). Notably, the inability of Nanog to rescue these genes further supports the notion that our SunTag cells have acquired LIF-independent self-renewal in the absence of functional Esrrb. In conclusion, these analyses underscore the complexity of the pluripotency network: whilst Nanog activates genes both in the presence and absence of LIF through regulatory regions belonging to clusters 1 to 4, at other enhancers the activation of Nanog can only be unmasked in the absence of LIF, when other pluripotency TFs are downregulated and their functional redundancy with Nanog, abolished. Unexpectedly, however, the genes repressed by Nanog in the absence of LIF, which exhibit a robust rescue of higher magnitude than that observed for the genes that Nanog activates (Fig. S4E), appear largely disconnected from the Nanog binding regions that we have studied here (Fig. 4F and Fig. S4F). These genes are associated, among other categories, with signalling and molecular pathways linked to differentiation (Fig. 4G). Hence, the probably indirect repression mediated by Nanog over these genes may underlie LIF-independent self-renewal in Dox-treated Sun-Tag cells, despite the lack of Esrrb and Klf4.

### Nanog sustains H3K27me3 to maintain silent differentiation genes upon LIF withdrawal

Gene set enrichment analysis (Fig. S5A) indicated that Nanog-rescued genes that are normally upregulated upon LIF withdrawal are enriched for targets of Polycomb Group proteins and for one of the marks they deposit to trigger facultative heterochromatin, H3K27me3 (FDR<6e-43). Hence, we profiled H3K27me3 in SunTag cells grown in the presence/absence of LIF and Dox. Overall, the patterns of H3K27me3 were found similar among all conditions, with notable exceptions (Fig. 5A). We identified three broad classes of H3K27me3 domains: those with high levels across all conditions and those that show either a loss or a gain of H3K27me3 upon LIF withdrawal (Fig. 5B and Table S3). Strikingly, the regions losing H3K27me3 in the absence of LIF maintained significant levels when endogenous Nanog expression was induced with Dox (Fig. 5B). In the presence of LIF, however, the induction of Nanog had minor consequences on H3K27me3, if any. This indicates that Nanog and LIF use parallel pathways to maintain H3K27me3 at a subset of H3K27me3 domains, and suggests that Nanog may confer LIF-independent self-renewal by sustaining H3K27me3 at these regions. Notably, the genes upregulated upon LIF withdrawal display differential enrichment among these three classes of H3K27me3 domains, depending on their Nanog-responsiveness: while nearly 70% of Nanog-rescued genes are marked by H3K27me3, which tends to decrease upon LIF-withdrawal except when Nanog is induced, only 30% of genes that are not rescued by Nanog show a similar pattern (Fig. S5B). This confirms that Nanog-rescued genes are particularly enriched in LIF-dependent H3K27me3. More specifically, H3K27me3 concentrates around the promoters of these genes, and displays a reduction in levels upon LIF withdrawal, exclusively in the absence of *Nanog* induction (Fig. 5C). Hence, Nanog stimulates the maintenance of H3K27me3 at a large subset of the promoters it represses in the absence of LIF. Nevertheless, a third of the genes repressed by Nanog in the absence of LIF is not embedded within H3K27me3; conversely, a third of the genes that are upregulated upon LIF withdrawal regardless of *Nanog* expression are enriched in H3K27me3 (Fig. S5B) and maintain higher levels around their promoters in the presence of Nanog (Fig. 5C). Thus, we determined whether quantitative differences regarding the effect of Nanog over these groups could be measured. Among the genes repressed by Nanog, the higher magnitude of rescue is observed for those genes that are embedded in H3K27me3 (Fig. S5C). Similarly, even though their gene expression changes upon Dox induction were not statistically significant, within the group of genes not rescued by Nanog, those enriched in H3K27me3 show a clear tendency to be downregulated (Fig. S5C). A clear, global pattern can be inferred from these analyses: the ability of Nanog to rescue genes that are upregulated upon LIF withdrawal is directly correlated with H3K27me3 levels. Accordingly, ordering the heatmap of genes upregulated upon LIF withdrawal (regardless of the ability of Nanog to rescue them) by their enrichment levels for H3K27me3 in the presence of LIF and the absence of Dox, naturally orders the genes from efficient to poor rescue (Fig. 5D). Hence, H3K27me3 levels before LIF withdrawal are highly predictive of the efficiency of Nanog to block gene upregulation during differentiation. Overall, these analyses indicate that in the absence of LIF, Nanog mediates its repressive function by other means than those described in its presence (Fig. 3): by maintaining high levels of H3K27me3.

**Fig. 5.**
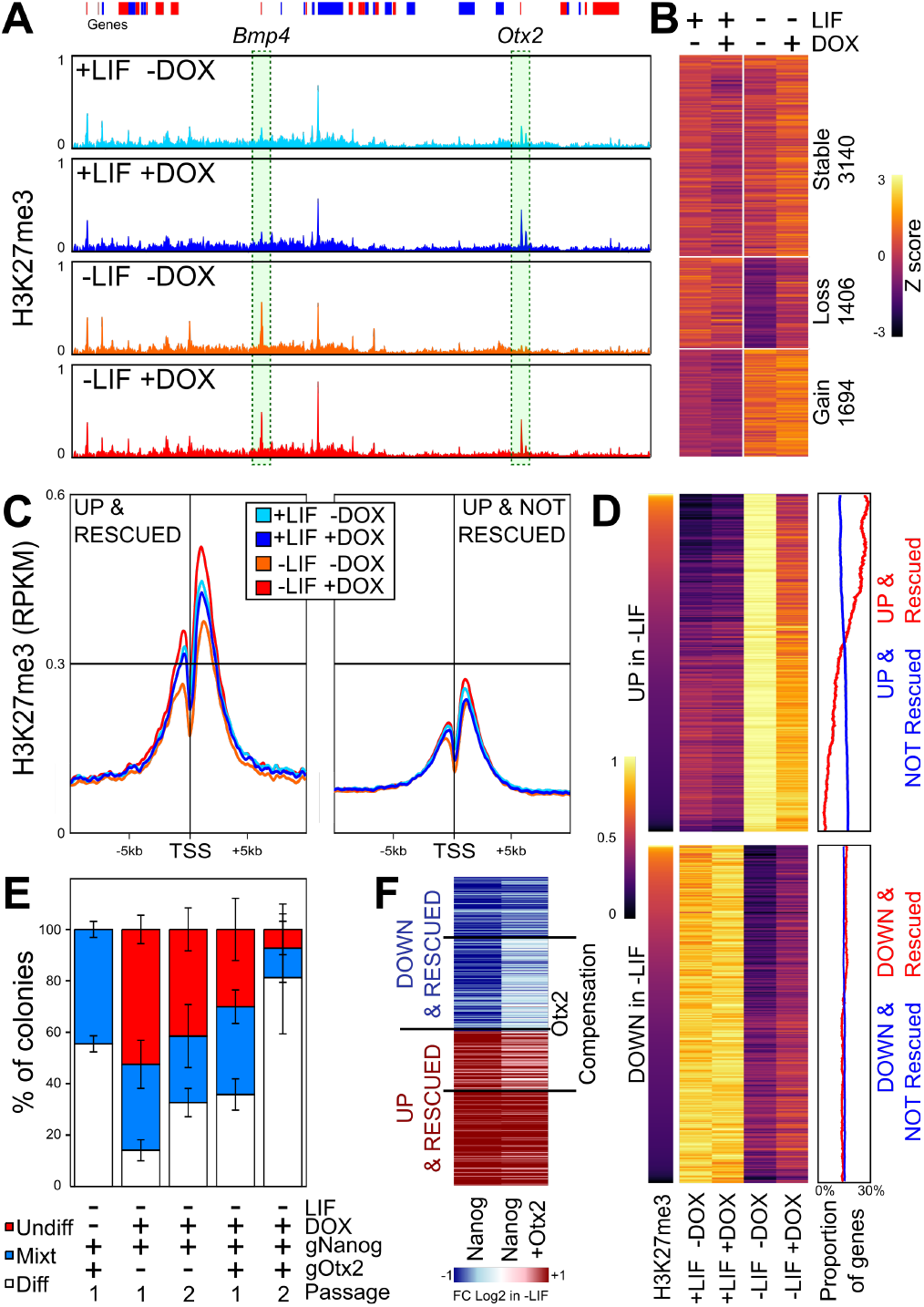
A *Nanog*/H3K27me3/*Otx2* axis controls LIF-independent self-renewal. **(A)** H3K27me3 average levels (reads per million) across 5.15Mb (mm9: chr14:45346123-50415313) encompassing the *Bmp4* and *Otx2* genes that display the two stereotypical behaviors observed in SunTag cells cultured in the presence/absence of LIF/Dox for 3 days. **(B)** Heat map of H3K27me3 z scores of 6,240 H3K27me3 domains identified in SunTag cells cultured as indicated. **(C)** Average H3K27me3 (reads per million) across promoter regions of genes upregulated in the absence of LIF and either rescued (left) or not (right) by Nanog. **(D)** Correlative analysis of ranked H3K27me3 (left) with gene expression changes (arbitrary units; middle) for transcripts upregulated (top) or downregulated (bottom) in Sun-Tag cells cultured in the presence/absence of LIF/Dox for 3 days. The percentage of genes belonging to rescued (red line) versus non-rescued genes (blue line) in a 500 gene sliding window across the regions of the heat map is shown on the right. **(E)** Histogram representing the percentage of undifferentiated (red), mixt (blue) and differentiated (white) colonies (Y-axis) counted after 6 (Passage 1) or 12 (Passage 2) days of clonal growth in the indicated conditions (X-axis). Error bars represent std. dev. (n=4). **(F)** Heat map of the average log2 fold change in gene expression between SunTag cells activating either *Nanog* or *Nanog* and *Otx2*, grown in the absence of LIF and the presence/absence of Dox for 3 days. Genes for which the simultaneous activation of *Otx2* compensates the changes observed when only *Nanog* is induced (FDR<0.05), are highlighted.

### Nanog represses *Otx2* to confer increased self-renewal efficiency regardless of LIF signalling

Among the genes downregulated by Nanog in the absence of LIF are several developmental TFs such as *Sox11*, *Id1* and *Pou3f1*, among others (Table S1). Therefore, it may be possible that LIF-independent self-renewal is attained by the simultaneous inhibition of several developmental pathways. However, within the list of Nanog-repressed genes characterised by Nanog-dependent H3K27me3, we could also identify *Otx2* (Fig. 5A and Table S3), a key regulator of the earliest stages of ES cell differentiation (**Acampora et al., 2013; Buecker et al., 2014**). Several lines of evidence point to *Otx2* down-regulation being an important mediator of Nanog function. First, *Otx2* has been already identified as an important negative target of Nanog (**Festuccia et al., 2012; Acampora et al., 2017**); accordingly, we observe its downregulation at the mRNA and protein levels upon *Nanog* induction (Fig. S5D, E). Moreover, further expression analyses indicate *Otx2* expression is closely controlled by Nanog levels (Fig. S5F). Second, the genes that are upregulated in the absence of LIF and that are rescued by Nanog, are enriched in genes activated by Otx2, while non-rescued genes are not (Fig. S5A). Third, the ectopic expression of Otx2 drives ES cells into differentiation (**Buecker et al., 2014; Acampora et al., 2017**), even in the presence of LIF, as we show here using our Sun-Tag system targeted to the *Otx2* promoter (Fig. S5E, G). Therefore, it may be possible that its Nanog-mediated down-regulation contributes to LIF-independent self-renewal in the context of our endogenous *Nanog* activation, despite the lack of strong upregulation of *Klf4* and *Esrrb*. To test this, we exploited the flexibility of the SunTag system to simultaneously activate *Nanog* and *Otx2* and perform clonal assays. Upon the additional induction of *Otx2*, the proportion of undifferentiated colonies decreased in the absence of LIF, compared to cells activating *Nanog* only, in particular after two successive rounds of clonal growth (Fig. 5E). These results clearly place *Otx2* as a key factor that needs to be repressed by Nanog in order to obtain efficient LIF-independent self-renewal. Next, we performed transcriptomic analyses upon *Nanog*/*Otx2* induction in the absence of LIF to identify the set of genes that were effectively compensated by the action of Otx2. We observed that around 40% of the genes repressed by Nanog, and 70% of the genes activated by Nanog, displayed similar levels to control cells grown in the absence of LIF and Dox, when both *Otx2* and *Nanog* were induced (Fig. 5F). Hence, at the molecular level, *Otx2* induction partially compensates the gene expression changes induced by Nanog overexpression in the absence of LIF, underscoring the antagonistic effect of Nanog and Otx2 over a large set of common genes (**Acampora et al., 2017**), including developmental TFs such as *Sox3*, *Sox11*, *Id1* and *Pou3f1*, among others (Table S1), which tend to be expressed in somatic cells and more particularly in neuroectodermal derivatives. In combination with the previous section, these results indicate strongly that Nanog controls H3K27me3 at key nodes in the differentiation network, such as *Otx2*, to indirectly repress a large set of genes involved with differentiation.

## Discussion

### Nanog rewires the pluripotency gene regulatory network

Gene regulatory networks constituted of master TFs are characterised by the capacity of individual factors to act over the same sets of regulatory elements, which together define and specify the molecular and transcriptional identity of each cell type (**Shlyueva et al., 2014; Spitz and Furlong, 2012**). However, we still have a relatively poor understanding of how single TFs impact globally on the recruitment of other members of a given network to impact its activity. Recently, the role of Oct4 has been suggested to rely on its ability to recruit Brg1 to render the chromatin accessible for other TFs to bind (**King and Klose, 2017**), matching a subset of the mechanisms we propose here for Nanog. However, it is unclear how much the initiation of differentiation that follows Oct4 depletion (**Niwa et al., 2000**) influenced the interpretations regarding how Oct4 directly impinges upon the pluripotency network. In our case, we have focused on Nanog, a factor that can be depleted from ES cells while preserving pluripotency (**Chambers et al., 2007**). Hence, it is likely that the rewiring of the network that we observe shortly after depleting Nanog, is due to primary and direct effects. Strikingly, our analyses suggest that the simplified view positing that pluripotency TFs bind cooperatively at regulatory elements to collectively control transcription, may need to be partially revisited: at least from the perspective of Nanog, the combinations of binding, their dependencies on Nanog, and their association to responsive genes, are more complex than we had previously anticipated (Fig. 6).

**Fig. 6.**
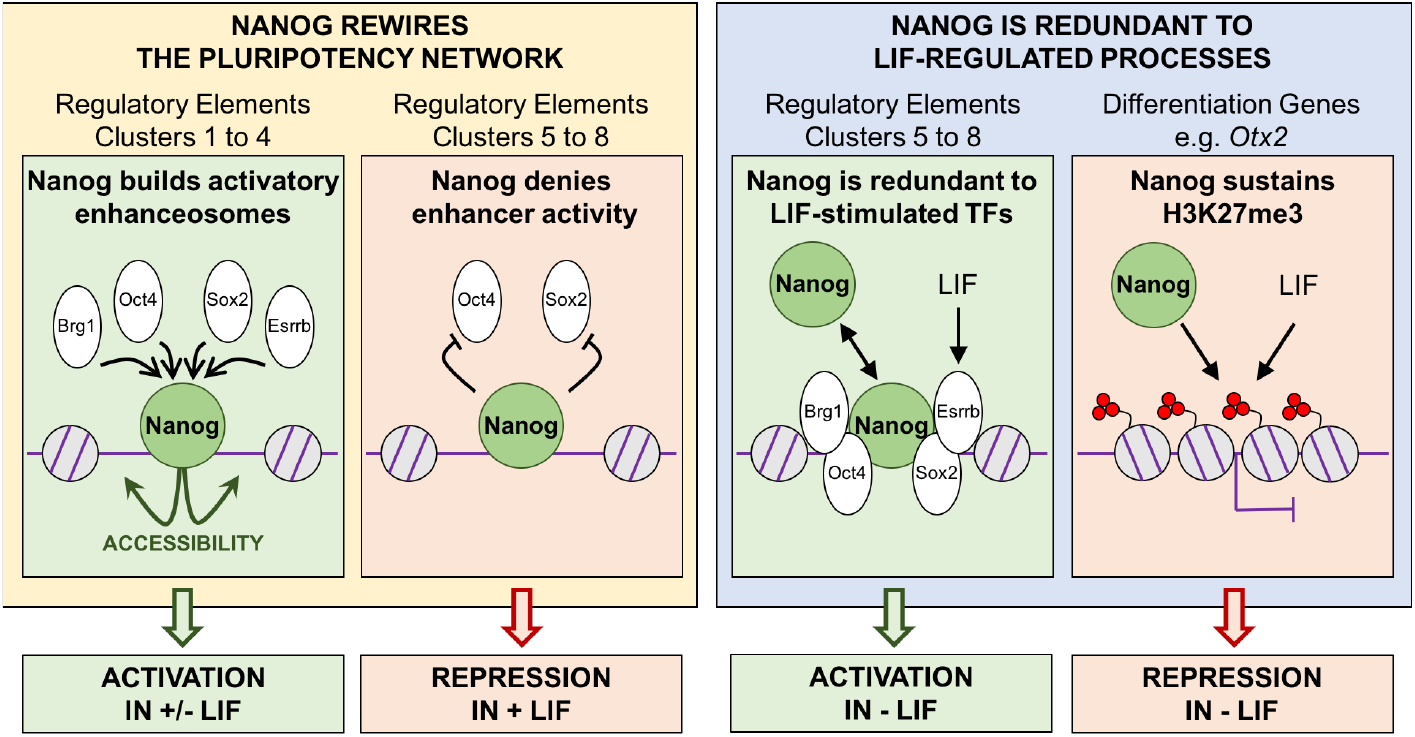
Nanog is a versatile TF impacting the pluripotency network and epigenome. The function of Nanog at stereotypical clusters of regulatory elements targeted by the pluripotency network, as well as at differentiation genes, is shown. Briefly, Nanog displays four major behaviours (left to right): 1/ recruitment of other factors (Oct4, Sox2 and Esrrb, together with Brg1) to promote chromatin accessibility and activate gene transcription; 2/ inhibiting enhancer activity, leading to gene repression either by blocking Oct4/Sox2 recruitment (shown) or by other mechanisms (not shown for simplicity; see text for details); 3/ complementing enhancer activity redundantly with other factors which are controlled by LIF (such as Esrrb) – in this case, its activatory role can only be appreciated in the absence of LIF; 4/ Nanog and LIF act in parallel to sustain H3K27me3 at differentiation genes such as Otx2. This latter role of Nanog is particularly important in the context of Nanog-mediated, LIF-independent self-renewal.

Although we observe, as expected, that Nanog-bound regions where other pluripotency TFs are also recruited are more strongly associated with Nanog-responsive genes compared to regions where Nanog binds alone, we also find that only a small subset displays similarly high binding levels of all three pluripotency TFs that we tested (Esrrb, Oct4 and Sox2). Indeed, the binding of one or two factors, in addition to Nanog, tends to dominate the others. This is valid even for Oct4/Sox2, which are believed to bind together at Oct4/Sox2 chimeric motifs (**Chew et al., 2005**). Therefore, different stoechiometries and/or residence times of individual factors seem to apply at distinct sets of regions. This produces a level of complexity that surpasses a simple model in which all factors bind together at key enhancers. Moreover, the effect of Nanog over these factors is also highly variable, with two clear groups of regions being easily identifiable: those in which Nanog plays a leading role; those where it does not. Strikingly, these two groups of regions display sharp distinctions regarding their association to Nanog-responsive genes. Regions where Nanog preserves chromatin accessibility and drives the recruitment of other pluripotency TFs and Brg1, are strongly biased to genes activated by Nanog. Conversely, the regions where Nanog does not promote TF binding or accessibility are associated either with genes repressed by Nanog in the presence of LIF, or with genes where Nanog acts as a redundant activator with LIF-stimulated TFs (Fig. 6).

To activate its targets, Nanog seems to use an expected mechanism essentially based on establishing a permissive chromatin architecture associated with the recruitment of other TFs and the formation of functional regulatory complexes. In contrast, while some precedents already pointed in the past into the direction of Oct4/Sox2-independent Nanog repression (**Navarro et al., 2008, 2012**), it is remarkable that Nanog-mediated repression is so strongly associated with regulatory elements where Nanog does not facilitate Esrrb, Oct4 and Sox2 binding and the chromatin remains equally accessible irrespectively of Nanog. This differential association between gene activation/repression and the role of Nanog as a nucleation factor of functional complexes, suggests that Nanog displays cooperative binding primarily to promote gene expression. At repressive sites, Nanog may either turn the whole complex into repressive or inhibit its otherwise transactivation potential. Additionally, at a large number of regions, the loss of Nanog leads to increased binding of either Oct4 or Sox2 (Fig. 6). It is therefore questionable that Nanog/Oct4 and Nanog/Sox2 bind at the same time over the same DNA molecules, at least over these regions. Thus, caution must be taken when extrapolating molecular functions from generic binding profiles: even though Nanog/Oct4 and Nanog/Sox2 appear to bind together, the binding of Nanog is in fact detrimental to that of the other two factors. Remarkably, these regions are closely associated with genes repressed by Nanog, indicating that to repress its targets Nanog interferes with the binding or the activity of other pluripotency TFs. Whether the alternate behaviours of Nanog to activate or repress transcription (Fig. 6) represents a general rule or a specific property of Nanog should be thoroughly investigated.

### Nanog-dependent H3K27me3 and *Otx2* repression, an alternative route for self-renewal

The ability of Nanog to block differentiation and promote LIF-independent self-renewal, something that not every pluripotent TF, including Oct4 and Sox2, is capable of doing, represents a defining property of Nanog (**Chambers et al., 2003**). However, this is not a unique characteristic of Nanog: a plethora of additional TFs, exemplified by Klf4 and Esrrb, have been progressively identified and demonstrated to provide LIF-independent self-renewal (**Niwa et al., 2009; Festuccia et al., 2012**). Hence, over the years, the importance of Nanog has been somehow equilibrated with that of other TFs, most notably Esrrb, which can replace Nanog in several contexts (**Festuccia et al., 2012; Zhang et al., 2018**). More strikingly, Esrrb has been proposed to be an obligatory mediator of the promotion of LIF-independent self-renewal by Nanog (**Festuccia et al., 2012**). Therefore, our observation of LIF-independent self-renewal in the absence of strong Esrrb upregulation, and of many other pluripotency TFs, is particularly enthralling. This is not the first time, however, that the role of a TF within the pluripotency network needs to be nuanced. Nanog itself was initially thought to be strictly required for germ cell development (**Chambers et al., 2007**) and for the induction of pluripotency via somatic cell reprogramming (**Silva et al., 2009**), conclusions that were subsequently invalidated (**Carter et al., 2014; Schwarz et al., 2014; Zhang et al., 2018**). Moreover, major experimental differences between our and previous studies may underlie the different conclusions regarding the mandatory requirement for Esrrb. Indeed, while others ectopically expressed Nanog constitutively and at high levels in *Esrrb* knock-out cells (**Festuccia et al., 2012**), we have used an inducible CRISPR-ON system to activate endogenous *Nanog* concomitantly with LIF withdrawal and the ensuing progressive downregulation of *Esrrb*. Therefore, the dynamic aspects of the two experimental setups are drastically different: it is possible that *Esrrb* is downregulated after Nanog has already impacted on other genes, which may independently block differentiation even when *Esrrb* is subsequently silenced. Besides this, however, our findings and their confrontation to previous conclusions highlight that different factors and mechanisms can potentially lead to similar phenotypic outcomes (**Konstantinides et al., 2018**).

Identifying the genes that mediate LIF-independent self-renewal in the absence of Esrrb may be particularly challenging because several prominent developmental regulators, from TFs to signalling molecules, are enriched among the genes that normally respond to LIF withdrawal but that endogenous *Nanog* induction is able to block. However, the genes that are upregulated when LIF is removed, and that Nanog is able to keep in check, display a blatant property: they tend to be targets of Polycomb PRC2 complexes and are embedded within H3K27me3 domains (**Azuara et al., 2006; Bernstein et al., 2006; Boyer et al., 2006**). At these genes, either LIF or Nanog are required to maintain H3K27me3 (Fig. 6). Therefore, this study identifies at least two modes of gene regulatory redundancy between Nanog and the LIF pathway: one directly based on LIF-stimulated TFs, such as Esrrb and Klf4, and another one based on the activity of Polycomb Group proteins (Fig. 6). We anticipate that identifying the exact molecular mechanisms used by Nanog to modulate H3K27me3 will be of great interest, and propose here that they are likely to be indirect. Overall, our observations argue for the existence of an alternative pathway to promote LIF-independent self-renewal through a previously unanticipated role of Nanog in the maintenance of H3K27me3 at differentiation-associated genes, thereby inhibiting the capacity of the cells to readily differentiate. This type of compensatory, chromatin-based mechanism, enables individual TFs to have broad impact by targeting key chromatin regulators with a more generic and systemic function. This remains so even when a regulatory network is largely dismantled, as is the case of the pluripotency network in the absence of LIF. This novel mechanism that we have unveiled may have the potential to dramatically increase the robustness and temporal integration of complex gene regulatory systems.

Whether the promotion of LIF-independent self-renewal associated with the inappropriate maintenance of H3K27me3 at differentiation genes results from the sum of many partial effects or, conversely, is based on specific and potent effects mediated by one or a few regulators, needs now specific attention. Given that the genes repressed by Nanog in the absence of LIF are strongly enriched for targets of Otx2, a driver of differentiation (**Acampora et al., 2013; Buecker et al., 2014**) that belongs to the category of bivalent genes in ES cells, we explored the possibility that *Otx2* downregulation may be the sole explanation for LIF-independent self-renewal in the absence of induced expression of other pluripotency TFs than Nanog itself. Using our SunTag cells to simultaneously activate both *Nanog* and *Otx2*, we could indeed observe that the efficiency of self-renewal was lower compared to the exclusive induction of *Nanog*. However, the effects became robust after two continuous phases of clonal growth, indicating that, for this level of upregulation of *Otx2*, the functional compensation takes time to be fully established. Nevertheless, our transcriptomic studies after three days of induction already show that a large fraction of the genes normally misregulated upon *Nanog* induction, show expression levels similar to control cells. In accord with the developmental role of Otx2 (**Acampora et al., 1995**), this seems to be particularly true for neuroectodermal genes. It should now be investigated whether a relationship similar to Nanog-Otx2 exists between Nanog and other lineage specific determinants targeted by H3K27me3. Overall, this work underlines the ability of *Nanog* to convey its function through remarkably distinct molecular mechanisms in different environmental conditions (Fig. 6). Extrapolating our work to other TFs of the pluripotency network and more generally to other gene regulatory systems, would be of great interest.

## Acknowledgements

We thank **Dr. Pentau Liu** for kindly providing PiggyBack plasmids and **Dr. Domingos Henrique** for fruitful discussions regarding the involvement of Polycomb activity in Nanog function. This work was supported by recurrent funding from the **Institut Pasteur**, the **CNRS**, and **Revive** (Investissement d’Avenir; ANR-10-LABX-73). P.N. acknowledges financial support from the **Fondation Schlumberger** (FRM FSER 2017), the **Fondation ARC** pour la recherche sur le cancer (ARC NAVARRO CA 14/12/2016), the **Agence Nationale de la Recherche** (ANR 16 CE12 0004 01 MITMAT), and the **Ligue contre le Cancer** (LNCC EL2018 NAVARRO). V.H is supported by the Fondation ARC and Revive. N.O. is supported by Revive.

## Author contributions

**V.H.** performed the experimental work with help from **I.G.** (ChIP and ATAC-seq), **F.M.** and **P.C.** (smFISH), **C.P.** (RNA-seq), **D.M.** (gRNA design) and **A.D.** (cell culture). **N.O.** analysed sequencing data. **V.H.**, **N.O.** and **P.N.** analysed and interpreted the results. **P.N.** designed the project and wrote the manusript with inputs from **V.H.** and **N.O.**

## SUPPLEMENTARY INFORMATION

Five supplementary figures accompany this manuscript:

**Fig.S1:** Characterisation of Dox-inducible SunTag ES cells.

**Fig.S2:** Validation of Nanog-responsive genes identified in this study.

**Fig.S3:** Heterogeneity of TF binding and chromatin accessibility throughout Nanog binding regions.

**Fig.S4:** Global and gene-specific responses of the transcriptome to Nanog induction in the absence of LIF.

**Fig.S5:** H3K27me3 and Otx2 are involved in LIF-independent self-renewal as established by Nanog.

They can be found at the end of this document.

Five Supplementary Tables are available online:

**Table S1:** The Nanog-centred transcriptome. In this table, all RNA-seq results are provided and annotated.

**Table S2:** Compendium of Nanog binding sites and their properties. In this table, all TF ChIP-seq and ATAC-seq results are provided and annotated.

**Table S3:** Nanog and LIF-dependent H3K27me3. In this table, all H3K27me3 domains are provided and annotated.

**Table S4:** Primer sequences and antibodies used in this study.

**Table S5:** Data Overview. In this table, all samples collected for RNA-seq, ChIP-seq and ATAC-seq are described along with Nanog public data sets.

**Supplementary Information, Fig. S 1.**
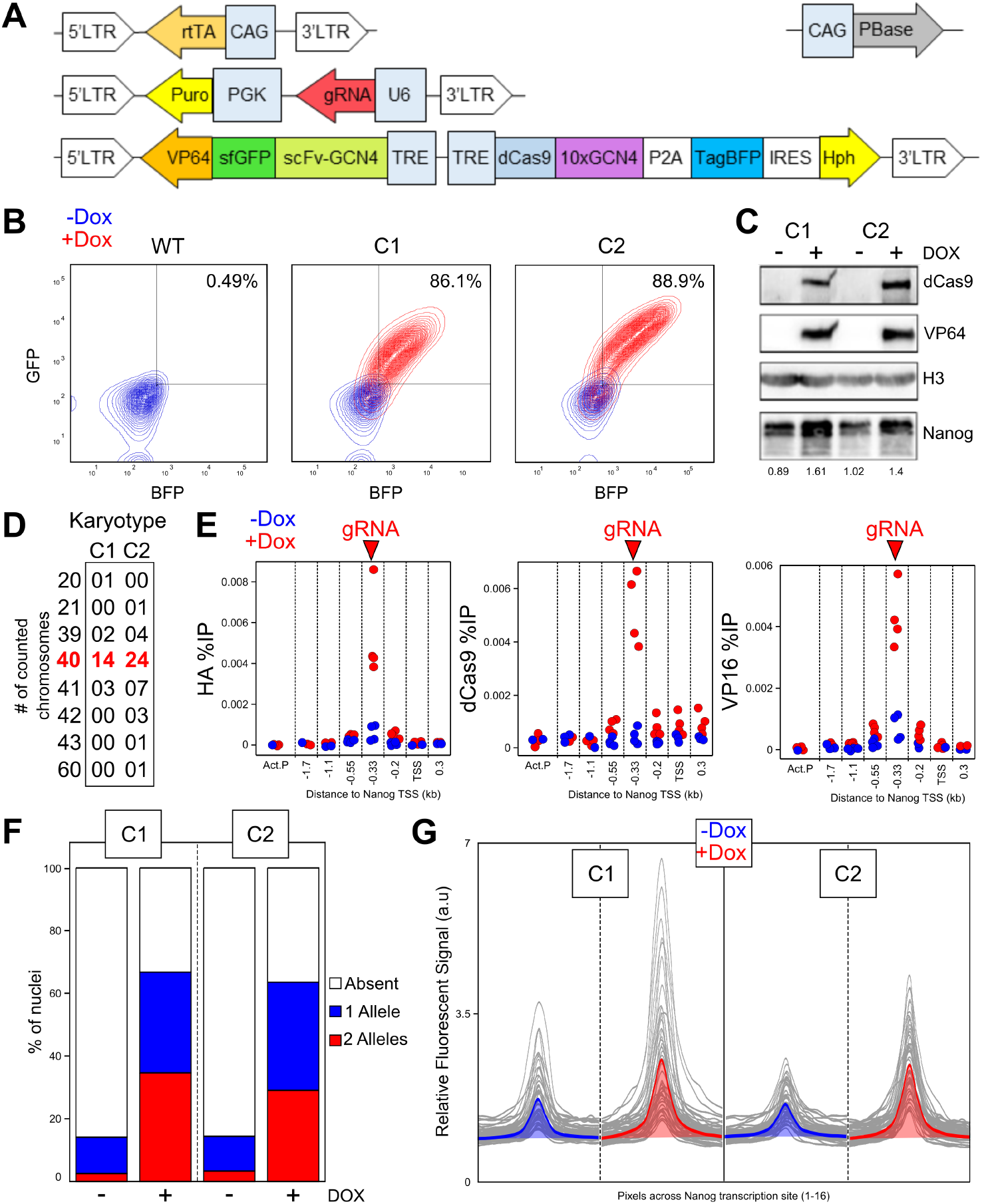
Characterisation of Dox-inducible SunTag ES cells. **(A)** Schematic representation of the Piggybac vectors used to generate ES cells expressing inducible SunTag transactivators. LTR: long terminal repeat; rtTA: reverse tetracycline-controlled transactivator; PBase: Piggybac transposase (non-integrative vector); CAG: constitutive RNAPII promoter; Puro: Puromycin resistance cassette; PgK: constitutive RNAPII promoter; U6: RNAPIII promoter for gRNA transcription; VP64: tetrameric fusion of Herpes simplex virus transactivation domain; sfGFP: super-folder green fluorescent protein; scFv-GCN4: Single-chain variable fragment antibody directed against the GCN4 yeast epitope; TRE: tetracycline responsive element; dCas9: enzymatically dead Cas9; 10xGCN4: 10 copies in tandem of the yeast GCN4 epitope; P2A: self-cleaving peptide; TagBFP: monomeric blue fluorescent protein; IRES: internal ribosome entry site; Hph: Hygromycin resistance cassette. **(B)** GFP versus BFP FACS profiles of wild-type cells (WT) and of two SunTag clones (C1 and C2) in the absence (blue) and the presence of Dox (red). The percentage indicates the proportion of double-positive cells. **(C)** Western-Blot analysis of the indicated proteins (right) in the indicated conditions (top). The numbers underneath indicate relative Nanog levels. **(D)** Karyotypes of SunTag ES cells. **(E)** ChIP analysis of the indicated proteins (the HA epitope is present in the two moieties of the SunTag system) at different positions along the *Nanog* locus (X-axis) with (red) and without Dox (blue). The amplicon overlapping the gRNA targeted sequence is indicated (top). Each data point represents an independent replicate (n=4; in both C1 and C2). **(F)** Proportion of SunTag cells displaying one or two *Nanog* active transcription sites as established by smFISH in the presence/absence of Dox for the two SunTag clones (n>500 nuclei in each clone/condition). **(G)** Relative quantification of the fluorescent signal measured along a line crossing Nanog transcription sites (n=40 sites in each clone/condition).

**Supplementary Information, Fig. S 2.**
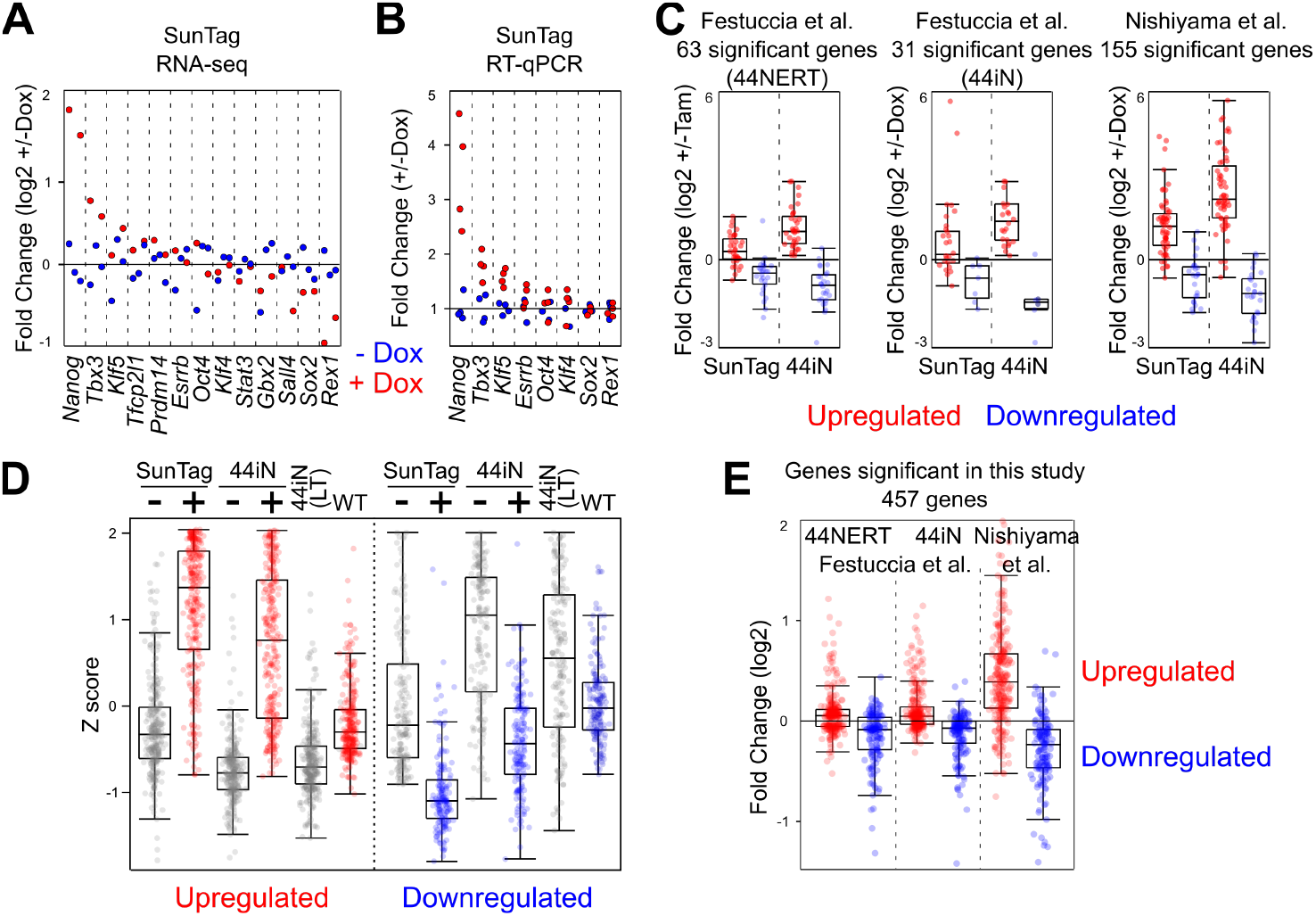
Validation of Nanog-responsive genes identified in this study. **(A)** Fold change (log2) of a set of pluripotency genes upon Dox treatment of SunTag cells. For each transcript, each value in the absence (blue) and in the presence (red) of Dox was normalised to the average of independent replicates. **(B)** RT-qPCR validation of a subset of the transcripts analysed in (A). **(C)** Confrontation of published results, as indicated, with our Nanog-responsive genes identified in SunTag and 44iN cells. **(D)** Box plot representation (z score, as in Fig. 2E) of expression levels of the genes identified in this study when considering both SunTag and 44iN together. 44iN(LT) indicate long-term culture in the absence of Dox (i.e. *Nanog* knock-out). **(E)** Quantification in published datasets, as indicated, of our extended list of Nanog-responsive genes.

**Supplementary Information, Fig. S 3.**
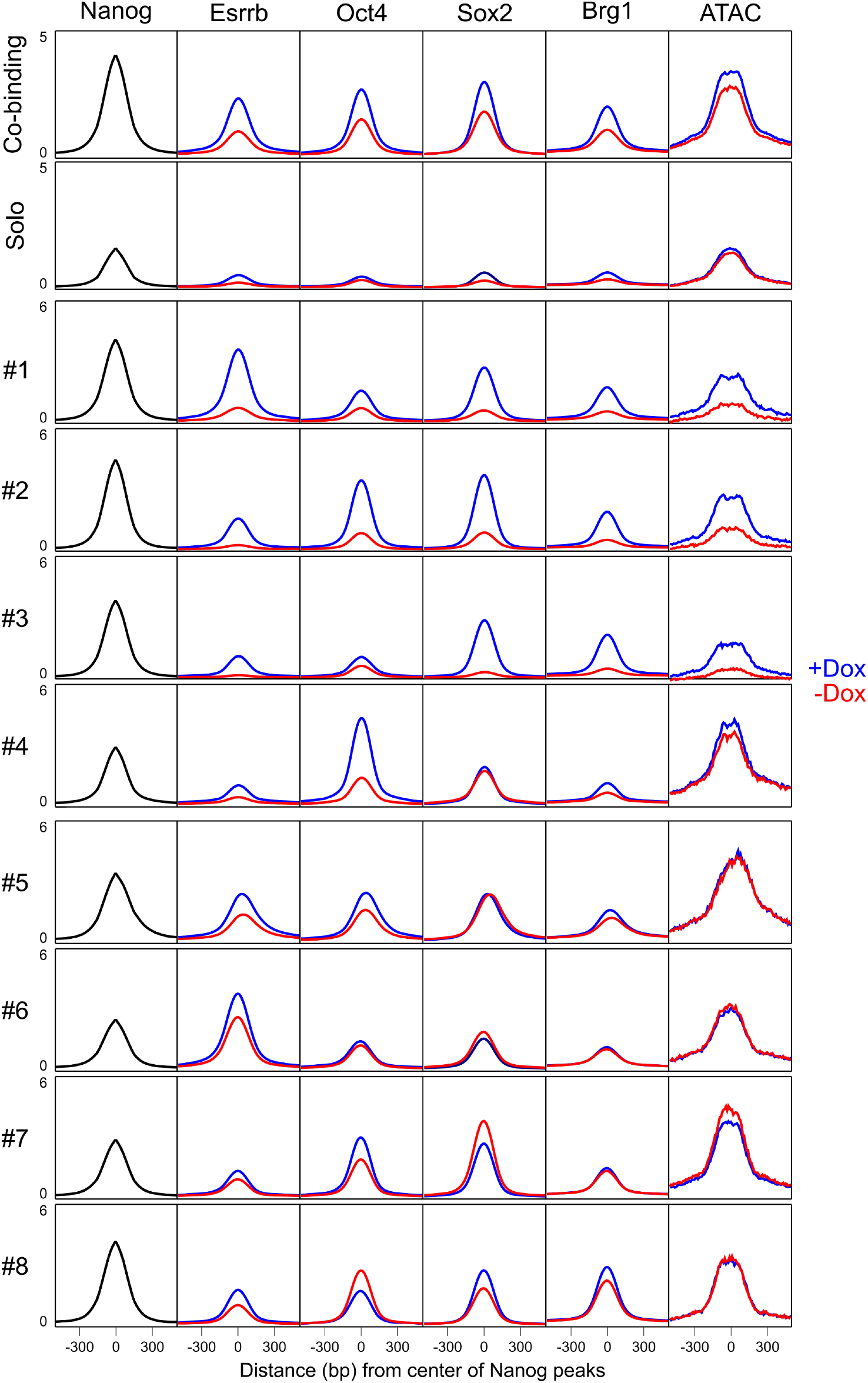
Heterogeneity of TF binding and chromatin accessibility throughout Nanog binding regions. The average binding profile of each factor (A.U. correcting for TF occupancy as described in Methods), as labelled on the top, is shown across Nanog peaks (summit at 0bp) for each Nanog subgroup identified in Fig. 3.

**Supplementary Information, Fig. S 4.**
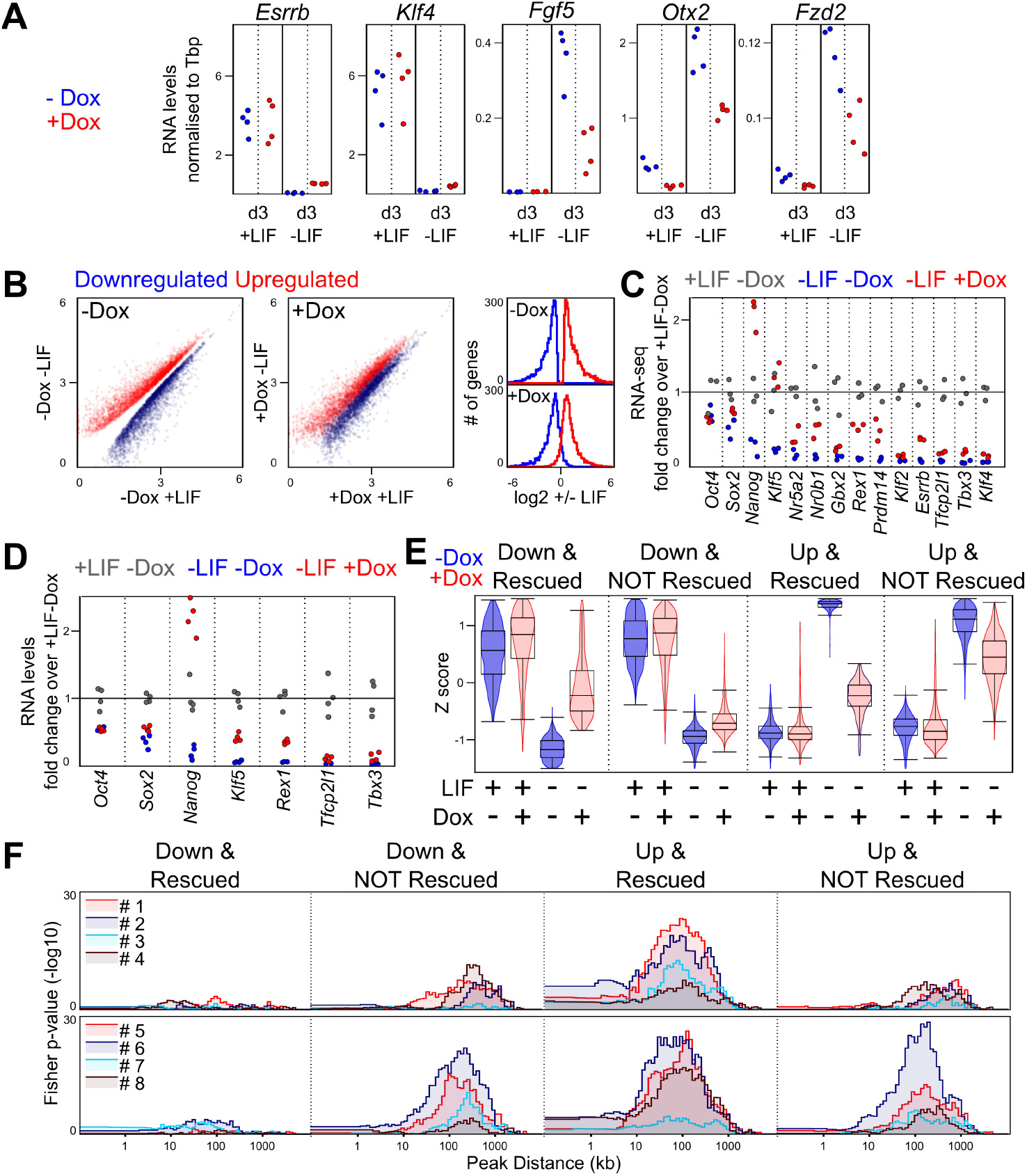
Global and gene-specific responses of the transcriptome to *Nanog* induction in the absence of LIF. **(A)** RT-qPCR of the indicated mRNAs across the conditions shown on the X-axis, in the absence (blue) and the presence (red) of Dox. Each data point represents individual measurements in both C1 and C2 clones (2 for each). **(B)** Left: scatter plot of normalised mRNA levels (DESeq2 normalised) of differentially expressed genes in the presence/absence of LIF, identified in the absence of Dox (FDR<0.05). Middle: Identical representation of the same set of transcripts but measured in the presence of Dox. Right: histogram showing the distribution of LIF-responsive genes across a range of fold changes (X-axis) in the absence (top) and the presence (bottom) of Dox. **(C)** Relative mRNA levels of a set of pluripotency factors normalised to the average expression of all replicates measured in the presence of LIF and the absence of Dox (n=3). **(D)** Validation by RT-qPCR of gene expression genes for a subset of pluripotency factors, presented like in (B). **(E)** Z score violin plots of the four groups of genes identified by RNA-seq in SunTag cells cultured in the absence/presence of LIF/Dox across the conditions indicated on the X-axis. **(F)** Association of the same four groups with the individual Nanog-binding clusters identified in Fig. 3, presented as in Fig. 3.

**Supplementary Information, Fig. S 5.**
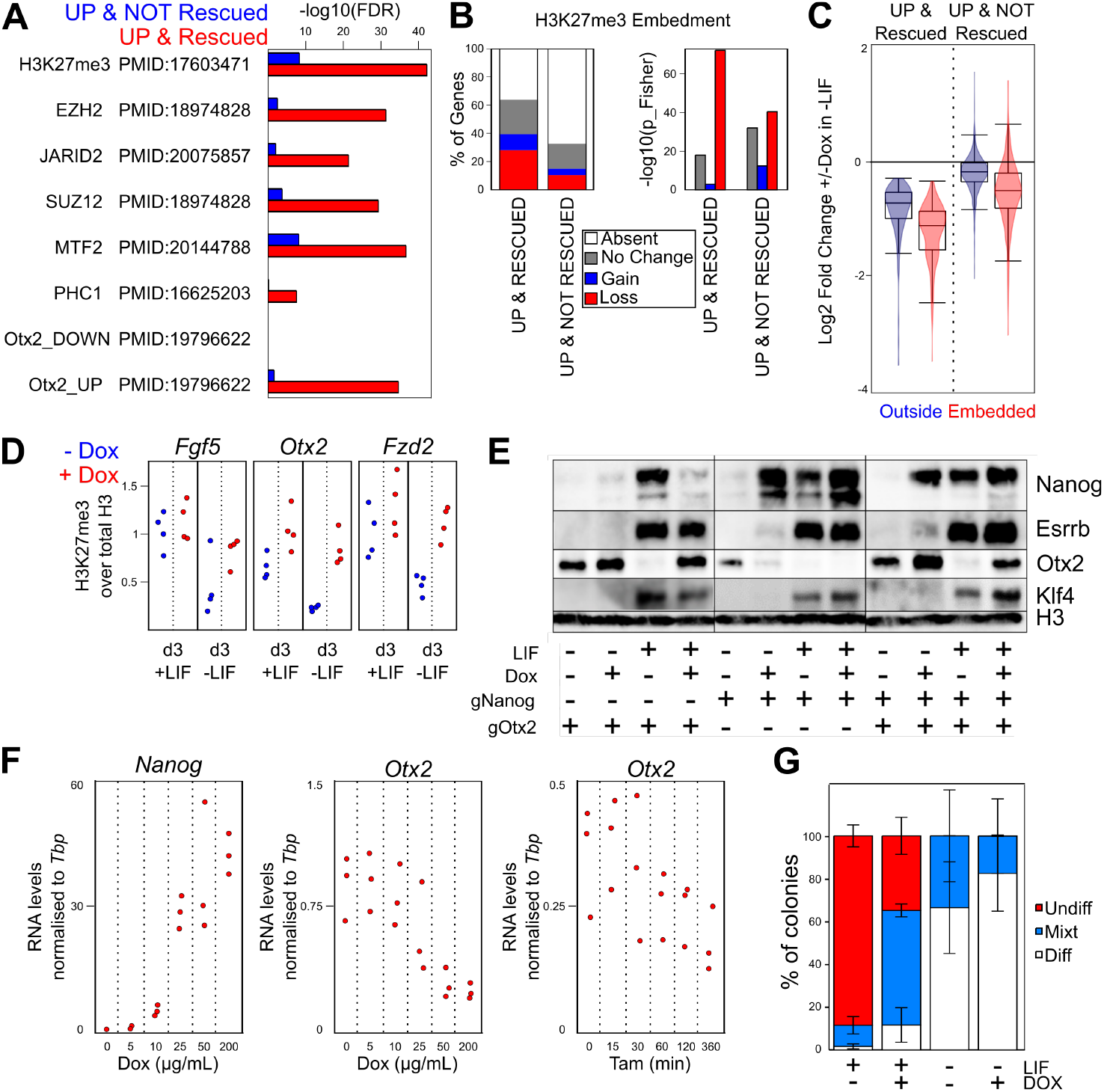
H3K27me3 and Otx2 are involved in LIF-independent self-renewal as established by Nanog. **(A)** Enrichment (-log10FDR) of UP and NOT rescued and UP and rescued genes for Polycomb Group targets and Otx2 responsive genes. The specific publication used by the Enrichr software (http://amp.pharm.mssm.edu/Enrichr/) to compute the enrichment is shown. **(B)** Proportion (left) and statistical significance (right) of H3K27me3-embedded genes activated without LIF and displaying differential rescue by Nanog (X-axis). **(C)** Violin plot of the log2 fold change of expression for the two previous categories of LIF-responsive genes split in function of their embedment (red) or not (blue) within H3K27me3 domains. **(D)** Normalised H3K27me3 ChIP-qPCR at three genes showing Nanog-dependent H3K27me3 enrichment in the absence of LIF. **(E)** Western Blot of the indicated factors (right) across multiple conditions (bottom). Note other WB presented in the principal figures correspond exactly to the blots shown here to facilitate direct comparisons between all analysed conditions. **(F)** Normalised mRNA levels of *Nanog* and *Otx2* over a Dox dose-response assay in 44iN and during a time-course of Tamoxifen treatment in Nanog-ERT2 fusion cells (44NERT), measured by RT-qPCR (n=3). 44NERT cells are *Nanog* knock-out cells expressing Nanog-ERT2 transgene (**Navarro et al., 2012**). (G) Clonal assay (n=4) in the indicated conditions (X-axis) using SunTag cells that activate *Otx2* exclusively.

